# PaperBLAST: Text-mining papers for information about homologs

**DOI:** 10.1101/133041

**Authors:** Morgan N. Price, Adam P. Arkin

## Abstract

Large-scale genome sequencing has identified millions of protein-coding genes whose function is unknown. Many of these proteins are similar to characterized proteins from other organisms, but much of this information is missing from annotation databases and is hidden in the scientific literature. To make this information accessible, PaperBLAST uses EuropePMC to search the full text of scientific articles for references to genes. PaperBLAST also takes advantage of curated resources that link protein sequences to scientific articles (Swiss-Prot, GeneRIF, and EcoCyc). PaperBLAST’s database includes over 700,000 scientific articles that mention over 400,000 different proteins. Given a protein of interest, PaperBLAST quickly finds similar proteins that are discussed in the literature and presents snippets of text from relevant articles or from the curators. PaperBLAST is available at http://papers.genomics.lbl.gov/.

## Introduction

Genome sequencing has accelerated the discovery of novel proteins far beyond the rate at which these proteins’ functions are being determined (Chang et al. 2016). Thus, to interpret genome sequences and to annotate the role of these predicted proteins, we rely on the similarity between the novel proteins and characterized ones. Proteins with over 30% similarity are likely to have the similar functions (Clark and Radivojac 2011), but to be 90% confident in the substrate of an enzyme, over 60% similarity may be required (Tian and Skolnick 2003).

Unfortunately, the databases of protein function that are used to make annotations are far from complete. As an example, the Swiss-Prot database (The UniProt Consortium 2017) is the largest curated resource of functional information about proteins, with experimental evidence relating to about 80,000 proteins. Nevertheless, the Swiss-Prot curators only curate 35-45% of new articles about protein function, and they focus on a few well-studied model organisms (Poux et al. 2016). They “do not have sufficient resources to actively curate organisms studied by smaller scientific communities” (ibid.).

As an alternative to expert curation, text-mining tools can find scientific articles that discuss a protein of interest (i.e., (Hoffmann and Valencia 2005)). With these tools, biologists can quickly find articles about a protein of interest and determine its function by reading the articles themselves, instead of relying on curators to do this for them. However, most of these text-mining tools have focused on model organisms and are not suitable for annotation by homology. Specifically, we are not aware of any text-mining tools that, given a protein of interest, search for information about similar proteins. There are tools that combine BLAST searches with links to articles from UniProt (Gilchrist et al. 2008) or from Genbank (Jaroszewski et al. 2014), but since these tools do not search the literature, their coverage is limited.

To allow access to the literature by homology, we developed the PaperBLAST web site (http://papers.genomics.lbl.gov/). Given a protein identifier or a protein sequence, PaperBLAST rapidly finds similar proteins that are discussed in the literature and provides links to those proteins and to articles about them.

## Results

### Examples

While we were studying the utilization of various carbon sources by *Pseudomonas fluorescens* FW300-N2E3, we discovered that the AO353_07705 protein is required for the utilization of L-carnitine (Price et al. 2016). As of April 2017, AO353_07705 is annotated in RefSeq (Tatusova et al. 2015) as “(Fe-S)-binding protein” (see WP_054594379.1) and is annotated by SEED (Overbeek et al. 2014) as “Predicted L-lactate dehydrogenase, Iron-sulfur cluster-binding subunit YkgF” (see FIG00138298). Neither of these annotations explained this protein’s role in carnitine utilization. Other annotation databases did not give useful results either: running InterProScan (Finn et al. 2017) or BLASTing against UniProt (The UniProt Consortium 2017) gave similarly vague information, and KEGG (Kanehisa et al. 2016) did not provide a prediction (see pba:PSEBR_a5225).

In contrast, PaperBLAST found published information about many homologs of AO353_07705, with the two closest homologs being from other strains of *Pseudomonas* (see Figure 1). This took less than 3 seconds. The articles about the closest homolog (from *P. syringae*) discuss gene regulation and might not be functionally informative. But one of the articles about the second homolog, PA5399 from *P. aeruginosa*, reports that it is required for the catabolism of glycine betaine and for the demethylation of dimethylglycine (Wargo et al. 2008). Although this level of detail is not apparent in PaperBLAST’s snippets, the snippets do mention a transposon mutant of PA5399, which makes it clear that of the five articles about the close homologs, this article is the most likely one to have functional information. Given the hypothesis that AO353_07705 is required for breaking down dimethylglycine, we can now explain the phenotype of the mutants in AO353_07705: carnitine is structurally related to glycine betaine, and some Pseudomonads break down carnitine via glycine betaine and dimethylglycine (Wargo and Hogan 2009).

**Figure 1:**
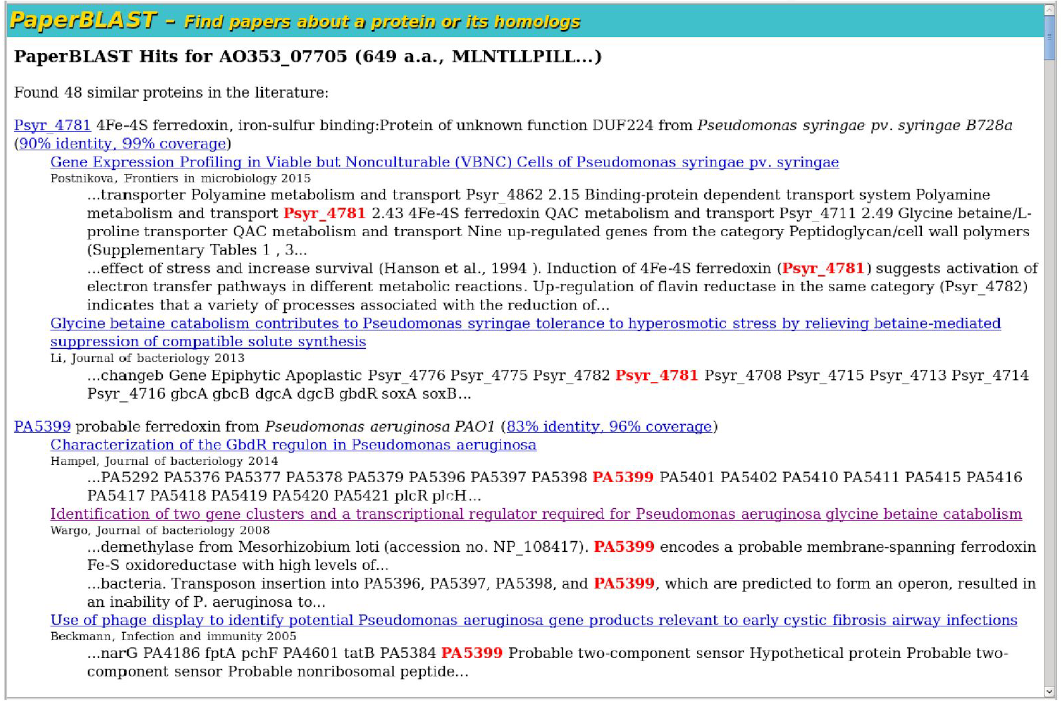
Example of PaperBLAST results. For each protein that is linked to the literature and is similar to the query, PaperBLAST shows a list of articles. For each article, PaperBLAST shows up to two snippets that mention the protein.

This example also illustrates that information about closely-related proteins may not propagate through the literature. Of the two articles that discuss the expression of the homolog from *P.syringae* (Psyr_4781), one article annotates the gene correctly (and cites (Wargo et al. 2008)). But the other article uses a vague description (“4Fe-4S ferredoxin”), even though it was published in 2015, seven years after the function of PA5399 was reported. This oversight reflects the difficulty of finding published information about similar proteins without an automated tool.

As another example, a colleague recently identified a phenotype for a mutant in PP_0995 from *Pseudomonas putida* KT2440 and asked us for advice as to its biochemical function. As of April 2017, the only relevant information in the protein family databases (according to InterProScan or the Conserved Domain Database (Marchler-Bauer et al. 2015)) was that PP_0995 belonged to an uncharacterized family (PF06532 or DUF1109). SEED annotated the protein as “extracytoplasmic function alternative sigma factor,” but we were not able to identify the rationale for this annotation. UniProt, RefSeq, and KEGG had uninformative annotations (accessions Q88P58, NP_743156.1, and ppu:PP_0995). In contrast, PaperBLAST identified nine homologs of this protein that are discussed in the literature, including two links to functional studies. The sixth hit (41% identity) was to CC3252 or NrsF from *Caulobacter crescentus*, which is proposed to be an anti-sigma factor: overexpression of NrsF prevents the induction of the SigF regulon after dichromate stress (Kohler et al. 2012). The ninth hit (33% identity) was to blr3039 or OsrA from *Bradyrhizobium japonicum*, which antagonizes the sigma factor EcfF and binds to it (Masloboeva et al. 2012). So we predict that PP_0995 is also an anti-sigma factor and that the SEED annotation is incorrect. Consistent with this, PP_0995 is adjacent to a putative sigma factor (PP_0994), as is common for anti-sigma factors. This example illustrates that an “uncharacterized” protein family may have functional information that has not been curated, that the rationale for a protein’s annotation is often unclear, and that PaperBLAST can be useful even if no close homologs of the query protein have been studied.

### The coverage of PaperBLAST

To assess the odds that PaperBLAST will provide useful information for a protein, we tested it on a random sample of predicted proteins (from sequenced genomes) that do not have specific functional annotations (see Methods). We did this test separately for proteins from bacteria, from plants, and from fungi. To describe the similarity of the best hit (if any) to the query, we used the ratio of the alignments’ bit score to the score for aligning the query against itself (the score ratio). Unlike the percent identity of the alignment, this takes the coverage of the alignment into account. To increase the odds that the two proteins have consistent domain content and function, we also required that the alignment cover at least 80% of the query. As shown in Figure 2, for 50% of bacterial proteins with vague annotations, PaperBLAST identified a homolog with a score ratio of over 0.4 that is discussed in the literature. These homologs are likely to have conserved functions (Clark and Radivojac 2011). For plant proteins and fungal proteins, the probability of finding a close homolog is a lower, but still reasonable (39% and 20%, respectively).

**Figure 2:**
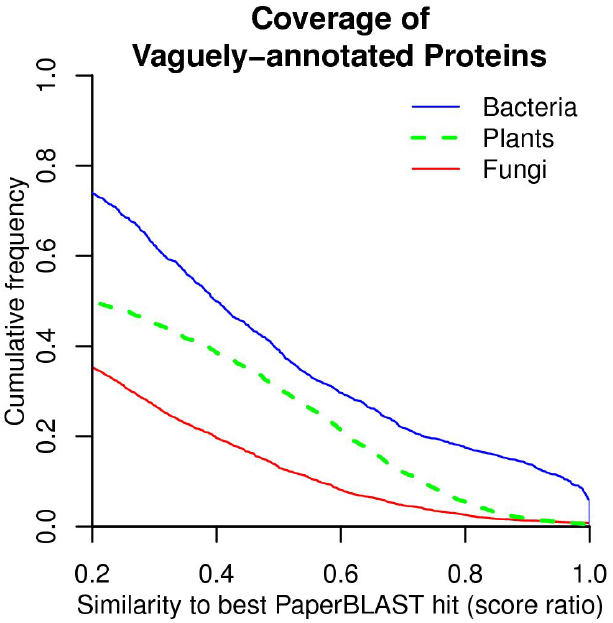
Coverage of PaperBLAST. We show how often hypothetical proteins, or other vaguely-annotated proteins, from different types of organisms have homologs in the PaperBLAST database with a BLAST score ratio above the given threshold. Only homologs with high-coverage alignments (at least 80%) were included.

However, just because a protein is discussed in the literature does not mean that there is any information about that protein’s function. So we checked a random sample of 20 of the vaguely-annotated bacterial proteins that have a close homolog (ratio above 0.4) in PaperBLAST. We found that in 12 of the 20 cases, PaperBLAST found a link to genetic or biochemical data about a close homolog’s function. In seven of the other eight cases, the article(s) discussed gene expression or protein expression data, which might not be informative as to the homolog’s function. In the remaining case, the article’s discussion of the homolog was based on the genome sequence only. So, we estimate that the odds of finding useful information in PaperBLAST about a vaguely-annotated bacterial protein is roughly 0.5 * 12/20 = 30%.

### PaperBLAST’s database of articles and proteins

PaperBLAST builds a database of protein sequences that are linked to scientific publications. Given this database and a protein sequence, PaperBLAST uses protein BLAST (Altschul et al. 1997) to find similar sequences. This only takes a few seconds.

PaperBLAST links proteins to articles in two ways: by searching the literature for mentions of protein identifiers by using the EuropePMC database (Europe PMC Consortium 2015), and by taking advantage of manually-curated resources (Swiss-Prot, GeneRIF, and EcoCyc). For an overview of how many different protein sequences are linked to articles by each resource, see Table 1. Overall, PaperBLAST contains 400,961 different protein sequences. If we cluster these sequences (Edgar 2010) at 80% identity, then there are 301,054 clusters. At 50% identity, there are 207,667 clusters.

**Table 1:** The number of proteins, scientific articles, and links between them in PaperBLAST’s database. Proteins with different identifiers but with the same sequence are only counted once. The total is less than the sum of the parts due to overlap between data sources. The count of proteins for Swiss-Prot includes some proteins that were linked to experimental evidence, but were not linked to articles about the protein’s function (see Methods). The count of proteins for EcoCyc does not include proteins that are not linked to any scientific articles (even though these are are included in PaperBLAST’s database).

| Source | #Proteins | #Papers | #Links | #Links with a snippet |
| --- | --- | --- | --- | --- |
| EuropePMC | 315,579 | 73,542 | 639,550 | 613,726 |
| Swiss-Prot | 79,388 | 27,453 | 38,342 | — |
| GeneRIF | 77,836 | 662,069 | 1,038,801 | — |
| EcoCyc | 3,923 | 11,143 | 22,769 | — |
| **Total** | 400,961 | 748,450 | 1,721,795 | — |

EuropePMC indexes the full text of 4.0 million articles and the abstracts of 26.9 million articles (http://europepmc.org/contentrss, accessed on February 23, 2017). PaperBLAST sends a query to EuropePMC for each protein identifier that appears in the open-access part of EuropePMC, in a PubMed abstract, or in the 499 most-referenced genomes. These identifiers are locus tags from MicrobesOnline (Dehal et al. 2010) or from RefSeq (i.e., b1234, ST1234, or ACICU_01235), RefSeq protein identifiers such as WP_000822979, or UniProt accessions such as P10018. We use queries of the form “identifier AND genus_name” to try to ensure that the article is actually discussing the protein as opposed to something else that happens to match a protein identifier. PaperBLAST also incorporates links between open access articles and UniProt or RefSeq identifiers from EuropePMC’s text mining efforts. Because EuropePMC uses different heuristics than we do, this yields additional links.

PaperBLAST also incorporates manually-curated information about protein function from Swiss-Prot, EcoCyc, and GeneRIF. First, PaperBLAST includes proteins in Swiss-Prot (the curated part of UniProt) if they have been studied experimentally. When a Swiss-Prot protein appears in the search results, PaperBLAST shows the manually-curated annotation as well as links to articles about the protein’s function, if these were provided by the curators. Second, PaperBLAST includes every protein from the well-studied bacterium *Escherichia coli*, as annotated in EcoCyc (Keseler et al. 2005). Again, PaperBLAST shows the manual annotation and a link to the articles selected by the curators. Finally, PaperBLAST incorporates the gene reference into function (GeneRIF) database from NCBI (Mitchell et al. 2003). GeneRIF links genes in Genbank or RefSeq to scientific articles in PubMed, and for each link, it also provides a short summary of the article’s claim about the gene’s function. For proteins that are linked to papers by GeneRIF, PaperBLAST shows the short summary.

For each automatically-generated link between a protein and an article in EuropePMC, PaperBLAST tries to select one or two snippets of text that include the protein identifier. 96% of these links have a snippet (Table 1). In contrast, PaperBLAST does not try to extract snippets from manually-curated articles: we expect that all of these articles are relevant and that the curator’s summary will be more useful than an automatically generated snippet.

### Completeness of PaperBLAST

Our impression is that EuropePMC’s coverage of recent articles about proteins is high, even for papers that are not open access, but we do not have a quantitative estimate. Some older articles are missing (for example, if the journal started depositing articles into PubMed Central in 2005, the older articles might be absent). EuropePMC still searches the abstract if it does not have access to the full text, but abstracts rarely include locus tags or other protein identifiers, so older articles are often missed. On the other hand, older articles are more likely to be curated.

To check the completeness of PaperBLAST, we examined experimentally-characterized proteins from two other resources: MetaCyc version 19.5 (Caspi et al. 2010) and the characterized proteins database CharProtDB (Madupu et al. 2012). (These are both curated resources; they date from 2015 and 2011, respectively.) We asked how often these proteins, or a nearly-identical homolog (with at least 99% identity and 90% coverage), appeared in the PaperBLAST database. We found that 75% of proteins in MetaCyc were already in PaperBLAST. Similarly, 70% of the proteins in the curated part of CharProtDB were already in PaperBLAST. (This is the portion entered by CharProtDB’s curators, as opposed to imported from Swiss-Prot or EcoCyc). This shows that PaperBLAST’s coverage is high, and it could be improved by incorporating additional curated resources (such as MetaCyc and CharProtDB). However the incremental benefit of adding each additional resource is modest: for example, MetaCyc contains 2,362 proteins that do not already have nearly-identical homologs in PaperBLAST.

## Discussion

Although PaperBLAST can often identify functional information for a homolog of a protein of interest, it is important to consider its limitations when using it.

First, as discussed above, many of the articles that mention a protein do not actually contain any information about that protein’s function. In particular, articles about gene expression often include tables of up-regulated or down-regulated genes. Because these articles mention many genes, they are over-represented in the PaperBLAST results. PaperBLAST’s snippets for these tables consist of locus tags, numbers, and gene annotations, rather than normal sentences, which is a hint that the article might not be useful. (Also, if you view these articles as a web page and search for the protein identifier, it will not appear, as the tables are rendered as images. You will need to view these articles as a PDF to find the mention of the gene.) In another common situation, the article mentions a protein that is believed to have a role in the pathway under consideration, but the experimental data concerns other proteins in the pathway. These articles can still be useful if they make a convincing prediction of the protein’s function. In the future, automated text analysis might be able to determine which articles have useful information about the protein’s function.

Second, PaperBLAST relies on protein identifiers, and it does not understand gene names like *yaaA*. This works well for non-model organisms, but some articles about proteins from model organisms are missed. The incorporation of manually-curated information from Swiss-Prot, GeneRIF, and EcoCyc will usually make up for this. A related issue is that PaperBLAST relies on matching by protein identifier. There are some spurious links due to other identifiers that happen to match protein identifiers. Spurious links should be obvious from the snippets, but more sophisticated text analysis might be able to filter them out. Also, many articles include DNA sequences such as primers, which could provide an independent check of what part of what genome is being studied (Haeussler et al. 2011).

Finally, a significant fraction of proteins are not similar to any protein that has been studied. For example, less than half of bacterial proteins have a homolog in PaperBLAST’s database with a BLAST score ratio of 0.3 or above. In the future, high-throughput approaches will provide functional information for proteins on a larger scale, and these data sets could be integrated into PaperBLAST. For example, Fitness BLAST (http://fit.genomics.lbl.gov/images/fitblast_example.html) identifies homologs that have mutant phenotypes in a collection of ∼4,000 genome-wide mutant assays from diverse bacteria (Price et al. 2016). Of the bacterial proteins that lack a reasonable homolog (ratio > 0.3) in PaperBLAST’s database, about 5% already have such a homolog with a mutant phenotype. To highlight these cases, a short summary of the Fitness BLAST results is shown at the bottom of the PaperBLAST results page.

In conclusion, PaperBLAST can quickly find articles that are relevant to the function of a protein of interest. We hope that it will be useful to every biologist who studies non-model organisms.

## Methods

### Searching EuropePMC

PaperBLAST uses 10 parallel threads to issue queries to EuropePMC. To avoid being blocked, each thread is limited to slightly less than one query every two seconds, or 4.5 queries per second in total. A recent build of PaperBLAST required 2.1 million queries, which takes roughly 500,000 seconds or 5 days. This is the slowest step in building the PaperBLAST database.

PaperBLAST also incorporates EuropePMC’s text-mined terms (ftp://ftp.ebi.ac.uk/pub/databases/pmc/TextMinedTerms). This gives us additional links from UniProt accessions and RefSeq accessions to scientific articles. For RefSeq, we incorporate only the mentions of protein entries.

### Selecting snippets of text from relevant articles

The main challenge in this step is to get access to the full text of the articles. PaperBLAST obtains the full text of open access articles as well as “author manuscript” articles from EuropePMC. Some other articles are available from the text-mining APIs of Elsevier (http://api.elsevier.com/documentation/FullTextRetrievalAPI.wadl) or CrossRef (http://tdmsupport.crossref.org/). Additional articles are downloaded using Maximilian Haeussler’s pubCrawl2 (https://github.com/maximilianh/pubMunch).

If the full text is available, then PaperBLAST identifies words that match the gene identifier. The gene identifier must appear as a separate word (possibly with punctuation at the end). For gene identifiers with a single letter followed by numbers, such as “c1406,” which might be more likely to have spurious hits, we also require that the case of the matches (i.e., “C1406” is ignored). If the full text is available but no snippet meeting these restrictions is found, then the link between the article and the gene is suppressed because it is probably a false positive. PaperBLAST records up to two snippets per gene per article, and each snippet is at most 160 characters, which may be considered fair use under copyright law.

If the full text is not available, PaperBLAST tries to use the abstract (from PubMed) to compute snippets.

### Information from manually-curated databases

PaperBLAST indexes proteins from Swiss-Prot (and shows them in the PaperBLAST results) if the Swiss-Prot curators identified experimental evidence about the protein. (This is indicated by the presence of the evidence code ECO:0000269 in the comment field.) However not all of these Swiss-Prot entries link to scientific articles in PubMed. A few of these entries are annotated based on theses or conference abstracts (or even direct submissions from scientists to UniProt). PaperBLAST does not link to these types of articles. A more common issue is that Swiss-Prot often links experimental evidence from a paper to the protein’s expression pattern or subcellular localization, which may not be functionally informative. PaperBLAST shows only the functional parts of the comment, and only links to the functionally-relevant articles. (Specifically, in the comment field, the “topic” that mentions the article must be one of function, catalytic activity, cofactor, enzyme regulation, disruption phenotype, or subunit structure.) After these restrictions, 48% of the Swiss-Prot entries with experimental evidence link to at least one article. Even if the article(s) are not shown on the PaperBLAST web site, the information is available via a link to the UniProt page for the protein.

PaperBLAST also indexes every protein from *Escherichia coli* K-12 in EcoCyc. Although most of the EcoCyc entries include links to articles in PubMed, about 200 do not -- these proteins will still appear in the PaperBLAST results if they are similar to the query.

To incorporate GeneRIF, PaperBLAST converts NCBI gene identifiers to RefSeq protein identifiers. To do this, it uses the NCBI Entrez programming utilities (https://eutils.ncbi.nlm.nih.gov/entrez/eutils/). About 84,000 gene identifiers were successfully converted to 78,000 distinct protein sequences.

### Redundant sequences

PaperBLAST uses usearch 8.0 (http://www.drive5.com/usearch/) to identify proteins in its database that have identical sequences. Duplicate proteins are removed from the BLAST database and are shown together in the output.

We also used usearch 8.0 (with the -cluster_fast and -sort length options) to determine the number of clusters of sequences at different levels of similarity.

### Settings for BLAST

PaperBLAST uses protein-protein BLAST 2.2.18 from NCBI’s BLAST+ package, with filtering of low-complexity sequences during lookup but not during alignment (the arguments -F “m S”). PaperBLAST reports hits with E < 0.001, up to a limit of 250 hits.

### Testing the coverage and completeness of PaperBLAST

To test the coverage of PaperBLAST on bacterial proteins, we used a randomly-selected set of 1,643 vaguely-annotated proteins from diverse bacteria that we described previously (Price et al. 2016). To obtain a random set of fungal proteins, we randomly selected 0.2% of the proteins in the fungal section of RefSeq (release 81, April 2017). We then selected the fungal proteins whose description matched “hypothetical,” “family,” or “domain”, which left a test set of 3,168 proteins. To obtain a random set of plant proteins, we randomly selected 1% of proteins from a random set of 20 genomes from Ensembl Plants release 34 (ftp://ftp.ensemblgenomes.org/pub/plants/release-34/fasta/). These genomes included five different species of the genus *Oryza*; the other 15 genomes were from unique genera. We then selected proteins whose description matched “Uncharacterized,” “hypothetical,” “family,” or “domain,” leaving a test set of 2,256 plant proteins.

All of these sequences were compared to the PaperBLAST database using protein-protein BLAST with E < 10^−5^, with filtering of low-complexity sequences during lookup but not during alignment. They were also compared to themselves to allow computation of the score ratios.

To test the completeness of PaperBLAST, we MetaCyc version 19.5 (from October 2015). In MetaCyc, the proteins.dat file links to 9,369 different UniProt entries. (There are also another 236 UniProt identifiers that are no longer current, which we ignored.) We also used CharProtDB, which we downloaded in April 2017 (but the most recent entry in CharProtDB was from 2011). In CharProtDB, the “curated” portion (which is unique to CharProtDB) contained 7,636 sequences. We compared these sequences to the PaperBLAST database using BLAST.

### Availability of code and data

The code for PaperBLAST is available at https://github.com/morgannprice/PaperBLAST.

All analyses were conducted with PaperBLAST database from April 2017, which is available at https://doi.org/10.6084/m9.figshare.4836407. It is based on searches against EuropePMC that were performed on or before March 26, 2017.

## Acknowledgements

We thank Johanna McEntyre and Jee-Hyub Kim for advice on using EuropePMC. We thank Maximilian Haeussler for providing pubCrawl2.

## Funding

This material by ENIGMA - Ecosystems and Networks Integrated with Genes and Molecular Assemblies (http://enigma.lbl.gov), a Scientific Focus Area Program at Lawrence Berkeley National Laboratory is based upon work supported by the U.S. Department of Energy, Office of Science, Office of Biological & Environmental Research under contract number DE-AC02-05CH11231.

